# Tfcp2l1 recruits Thrap3 to sustain ESC self-renewal via *Jmjd1c* exon-5 splicing and DNA repair choice

**DOI:** 10.64898/2026.11.05.764592

**Authors:** Wei Lu, Xiaoxiao Wang, Yuexuan Li, Peng Chen, Yuqing Zhu, Shou-Dong Ye

## Abstract

How embryonic stem cells (ESCs) maintain genomic integrity while sustaining self-renewal remains poorly understood. We find that transcription factor CP2-like 1 (Tfcp2l1) recruits Thrap3 in mouse ESCs, where this complex maintains RNA polymerase II activity and promotes transcription of Tfcp2l1 target genes. In addition, *Thrap3* supports mESC self-renewal. Interestingly, knockdown of either *Tfcp2l1* or *Thrap3* induces highly overlapping alternative-splicing changes, including reduced inclusion of exon-5 in *Jmjd1c*. Further studies reveal that skipping of *Jmjd1c* exon-5 weakens the association between Jmjd1c and Mdc1, accompanied by elevated Mdc1-associated dimethylation, leading to reduced Mdc1 and Rnf8 interaction and diminished K63-linked ubiquitin signalling. These alterations impair chromatin recruitment of the Rap80 and Brca1 repair complex and increase Rad51 foci formation, indicative of a shift in DNA repair pathway preference. Accordingly, loss of Tfcp2l1 or Thrap3 results in accumulated DNA damage, hypersensitivity to genotoxic stress, compromised ESC self-renewal and enhanced apoptosis. Similar phenotypes can be recapitulated in human ESCs. Together, our findings provide new insights into how ESCs coordinate genome maintenance with self-renewal.

## 1. Introduction

Mouse embryonic stem cells (mESCs) are defined by their sustained self-renewal and their pluripotency to generate derivatives of all embryonic lineages (Byers *et al*, 2022; Nichols & Smith, 2009). Their rapid proliferation and distinctive cell-cycle organization expose them to substantial replication-associated stress, making genomic integrity essential for maintaining stem-cell identity and developmental potential (Blakemore *et al*, 2021; Weissbein *et al*, 2014). Because mESCs have a permissive G1/S checkpoint and progress rapidly through the cell cycle (Liu *et al*, 2019; ter Huurne *et al*, 2017), they must coordinate DNA replication with effective surveillance and repair mechanisms to limit the accumulation of genomic lesions. This challenge is compounded by distinctive replication dynamics, including a high density of replication origins and relatively slow replication-fork progression, which can generate replication stress (Jodkowska *et al*, 2022; Kurashima *et al*, 2024). MYBL2 and ATM have been implicated in limiting this stress in ESCs (Blakemore *et al*., 2021). To maintain genome stability, ESCs rely on Atm- and Atr-dependent DNA damage response (DDR) pathways (Roy *et al*, 2025), while the elimination of irreparably damaged cells through apoptosis helps prevent the propagation of genomic abnormalities (Huskey *et al*, 2015). ESCs have also been reported to favour homologous recombination (HR) over classical non-homologous end joining (NHEJ), even outside S phase (Merkle *et al*, 2017; Serrano *et al*, 2011), reflecting their distinctive requirements for genome maintenance. Thus, genomic integrity is a prerequisite for sustained self-renewal rather than merely a consequence of it. However, the balance between DNA repair and cell-fate control is critical (Chen *et al*, 2021; Weissbein *et al*., 2014). Insufficient repair allows genomic lesions to accumulate, while dysregulated DDR signalling can trigger cell death or impair pluripotency (Merkle *et al*, 2022). Human pluripotent stem cells are particularly susceptible to such risks, as genetic alterations including TP53 mutations and chromosomal abnormalities may be selected for during prolonged in vitro culture (Benvenisty *et al*, 2025; Li *et al*, 2026). It remains incompletely understood how the core pluripotency circuitry participates in genome maintenance and shapes DNA damage response regulation (Dong *et al*, 2026; Tang *et al*, 2026).

Transcription factor CP2-like 1 (Tfcp2l1) is a well-established regulator of ESC self-renewal and pluripotency (Ye *et al*, 2013; Zhang *et al*, 2021). It acts downstream of the LIF/Stat3 and Wnt/β-catenin signalling pathways (Martello *et al*, 2013; Qiu *et al*, 2015), and its ectopic expression can support the maintenance of an undifferentiated state and promote the reprogramming of post-implantation epiblast stem cells (Hancock *et al*, 2021; Sun *et al*, 2018). In addition to its transcriptional functions, the N-terminal region of Tfcp2l1 mediates interactions with regulatory partners, as exemplified by its association with *Mta1* in the control of lineage commitment (Liu *et al*, 2017). This finding suggests that the N-terminal region may serve as a platform for interactions with distinct co-regulatory factors, raising the possibility that Tfcp2l1 influences cellular processes beyond direct transcriptional regulation. One potential connection is the coupling between transcription and pre-mRNA splicing. RNA polymerase II (Pol II) elongation and phosphorylation of its C-terminal domain (CTD), particularly at Ser2, are associated with co-transcriptional RNA processing (Appel *et al*, 2023; Egloff & Murphy, 2008; Jonkers & Lis, 2015), providing a potential mechanism through which transcriptional regulation can influence alternative splicing (AS). Notably, Tfcp2l1 exerts opposing effects across cancers, acting as an oncogene in bladder (Heo *et al*, 2022) and hepatocellular (Qiu *et al*, 2024) carcinoma but as a tumour suppressor in clear cell breast (Wang *et al*, 2026) and thyroid (Zeng *et al*, 2024) carcinoma, indicating context-dependent outcomes. Given the distinctive replication-associated challenges faced by ESCs, it is important to determine whether Tfcp2l1 engages RNA-processing co-factors through its N-terminal docking region, and whether such interactions contribute to genome maintenance in ESCs. This possibility has not been examined.

Here, we identify a functional link between the pluripotency-associated factor Tfcp2l1 and DNA damage signalling in ESCs. Tfcp2l1 interacts with the RNA-processing factor Thrap3, and Tfcp2l1-dependent Thrap3 recruitment alters RNA Pol II regulation and AS of *Jmjd1c* exon-5. Changed Jmjd1c exon-5 inclusion remodels Mdc1-associated dimethyl-lysine signals, disrupts Mdc1 and Rnf8 interaction, and impairs recruitment of downstream repair factors including Rap80 and Brca1. Disruption of this regulatory network increases DNA damage and compromises ESC maintenance, with equivalent phenotypes observed in human ESCs. Together, these findings indicate how pluripotency-associated transcription sustains genome integrity in ESCs.

## 2. Results

### 2.1. Thrap3 is a direct interacting partner of Tfcp2l1 in mESCs

To identify proteins binding to Tfcp2l1, we expressed FLAG-tagged mouse *Tfcp2l1* in mESCs cultured in the presence of LIF (EV-*Tfcp2l1*), using empty vector (EV)-transfected cells as the control, and performed three independent IP-LC-MS/MS experiments (Fig. 1A; Fig. S1A). Of 137 proteins identified in the Tfcp2l1 immunoprecipitate, 26 were annotated as transcription-associated and 16 as related to ESC self-renewal; intersection analysis identified 5 proteins with both functions (Sequest HT score > 5): *Ddx3x*, *Hnrnpk*, *Thrap3*, *Hnrnpa2b1* and *Hnrnph1* (Fig. 1B,C; Supplementary Table 1). Whereas *Ddx3x*, *Hnrnpk*, *Hnrnpa2b1* and *Hnrnph1* are well characterized in ESCs, *Thrap3* has received only limited attention, so we verified this interaction biochemically. In endogenous co-immunoprecipitation (Co-IP) assays, reciprocal pulldowns using anti-Thrap3 and anti-Tfcp2l1 antibodies recovered the respective protein complexes from mESCs (Fig. 1D,E). We also constructed HA-tagged mouse *Thrap3* (EV-*Thrap3*) and verified the transfection efficiency (Fig. S1B,C). Exogenous Co-IP assays in mESCs transfected with FLAG-*Tfcp2l1* recovered Thrap3, and parallel exogenous Co-IP assays in mESCs transfected with HA-*Thrap3* recovered Tfcp2l1 (Fig. S1D,E). Furthermore, immunofluorescence showed nuclear co-localization with overlapping double-channel intensity profiles (Fig. 1F; Fig. S1F). Thus, *Tfcp2l1* directly interacts with *Thrap3* in mESCs.

**Figure 1.**
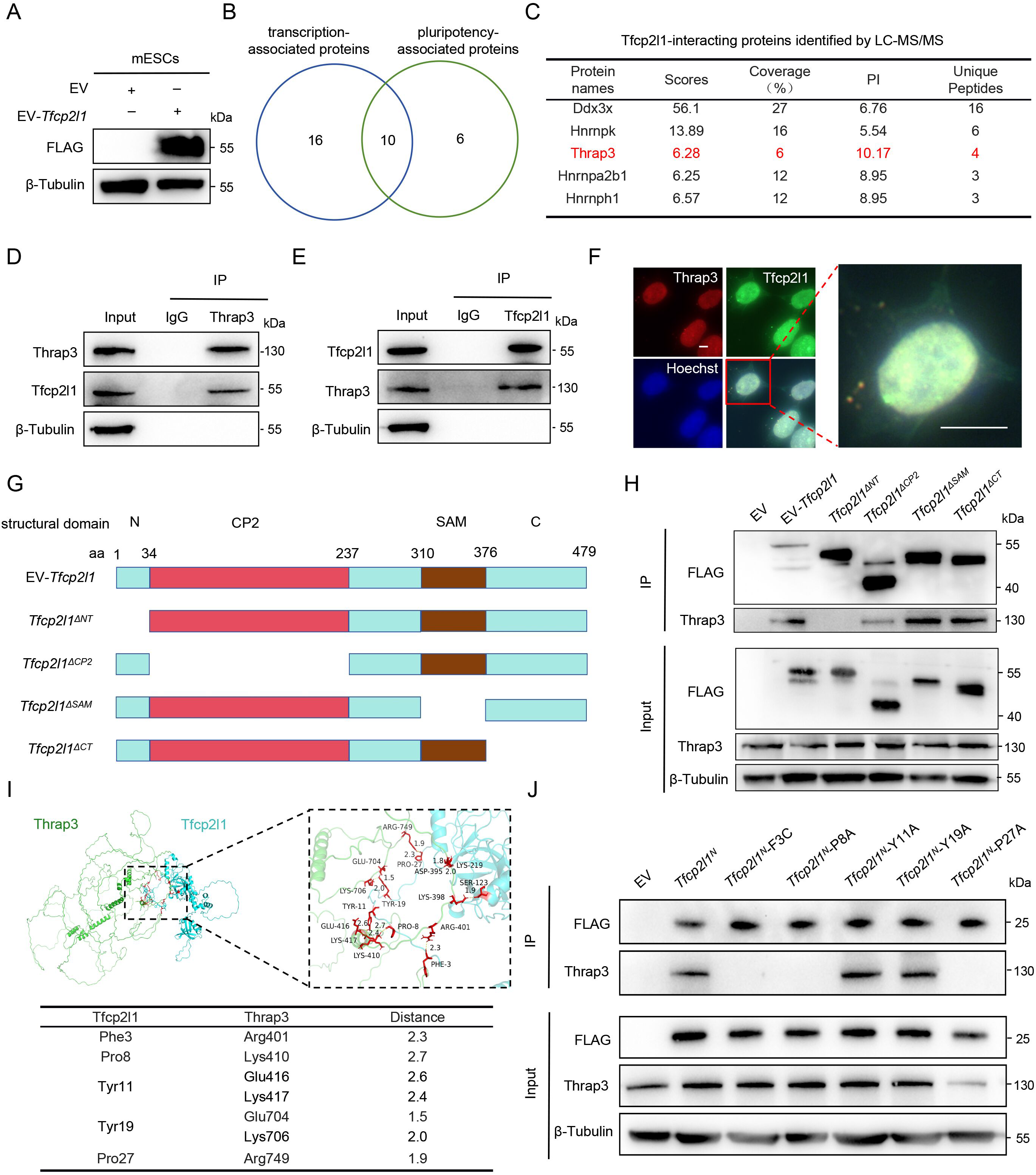
Thrap3 binds to the N-terminal domain of Tfcp2l1 in mESCs. **A** Western blot analysis of mESCs transfected with FLAG-tagged mouse *Tfcp2l1* (EV*-Tfcp2l1*) or empty vector (EV). **B** Intersection analysis was conducted for 26 transcription-associated proteins and 16 pluripotency-associated proteins from immunoprecipitated complexes captured by three independent FLAG IP-LC-MS/MS experiments. **C** The top 5 highly enriched proteins (enrichment score > 5) were selected from the intersecting proteins in (B). **D**, **E** Endogenous Co-IP assesses the interaction between Tfcp2l1 and Thrap3. Immunoprecipitation was performed using anti-Thrap3 antibody (D) and anti-Tfcp2l1 antibody (E), respectively. **F** Immunofluorescence staining of the Thrap3 (red) and the Tfcp2l1 (green) in mESCs. Scale bar, 5 μm. **G** Schematic of plasmid constructs encoding *Tfcp2l1* with different domain truncations. **H** Immunoprecipitation analysis of the enrichment of Thrap3 in mESCs after transfection with different FLAG-tagged truncated *Tfcp2l1* plasmids. **I** The three-dimensional structure model of the Thrap3 and Tfcp2l1 complex was constructed using molecular docking. The specific binding sites in the N-terminal regions of the two proteins were predicted, and the structure was visualized using PyMOL. **J** Immunoprecipitation analysis of the enrichment of Thrap3 in mESCs transfected with FLAG-tagged *Tfcp2l1^N^* and its N-terminal F3C, P8A, Y11A, Y19A and P27A single-point mutant plasmids.

To map the interaction interface, we generated a series of truncation constructs. Truncation mapping localized the interaction to the N-terminus of Tfcp2l1 (aa 1-34) (Fig. 1G,H) and to the Thrap3/Bclaf1 family domain of Thrap3 (aa 191-766) (Fig. S1G,H). Subsequently, we built a complex model via HADDOCK docking and visualized the structure in PyMOL to predict the putative contacting amino acid residues (Fig. 1I). Single-point mutagenesis further revealed that Phe3, Pro8 and Pro27 of Tfcp2l1 together with Arg401, Lys410 and Arg749 of Thrap3 were required for their physical interaction (Fig. 1J; Fig. S1I), indicating that the N-terminal segment of Tfcp2l1 constitutes the docking platform for its Thrap3 partner.

### 2.2. Thrap3 promotes the maintenance of mESC stemness

To investigate the role of Thrap3 in the self-renewal of mESCs, we cultured EV and EV-*Thrap3* mESCs in the presence of LIF for 7 days. Compared with EV control cells, *Thrap3*-overexpressing cells exhibited increased protein levels of the self-renewal markers Klf4 and Oct4, while the levels of the differentiation markers Gata4 and Gata6 were decreased (Fig. 2A). In addition, overexpression of *Thrap3* enhanced the colony-forming capacity and alkaline phosphatase (AP) activity of mESCs following 7 days of LIF withdrawal (Fig. 2B). RT-qPCR revealed increased mRNA expression of *Nanog*, *Sox2*, *Oct4* and *Klf4*, alongside reduced transcript levels of *Cdx2*, *Oct6* and *Gata4* in *Thrap3*-overexpressing mESCs cultured in the presence of LIF for 7 days (Fig. 2C). Moreover, Sox2 fluorescence intensity was elevated in *Thrap3*-overexpressing cells (Fig. 2D). During embryoid body (EB) differentiation, *Thrap3* overexpression sustained *Oct4* and *Sox2* expression for longer relative to EV cells, and the induction of lineage markers *Elf5*, *Gata4*, *Gata6*, *Cdx2*, *Sox17* and *T* was attenuated (Fig. 2E; Fig. S1J,K). These findings indicate that *Thrap3* overexpression promotes maintenance of the undifferentiated state of mESCs.

**Figure 2.**
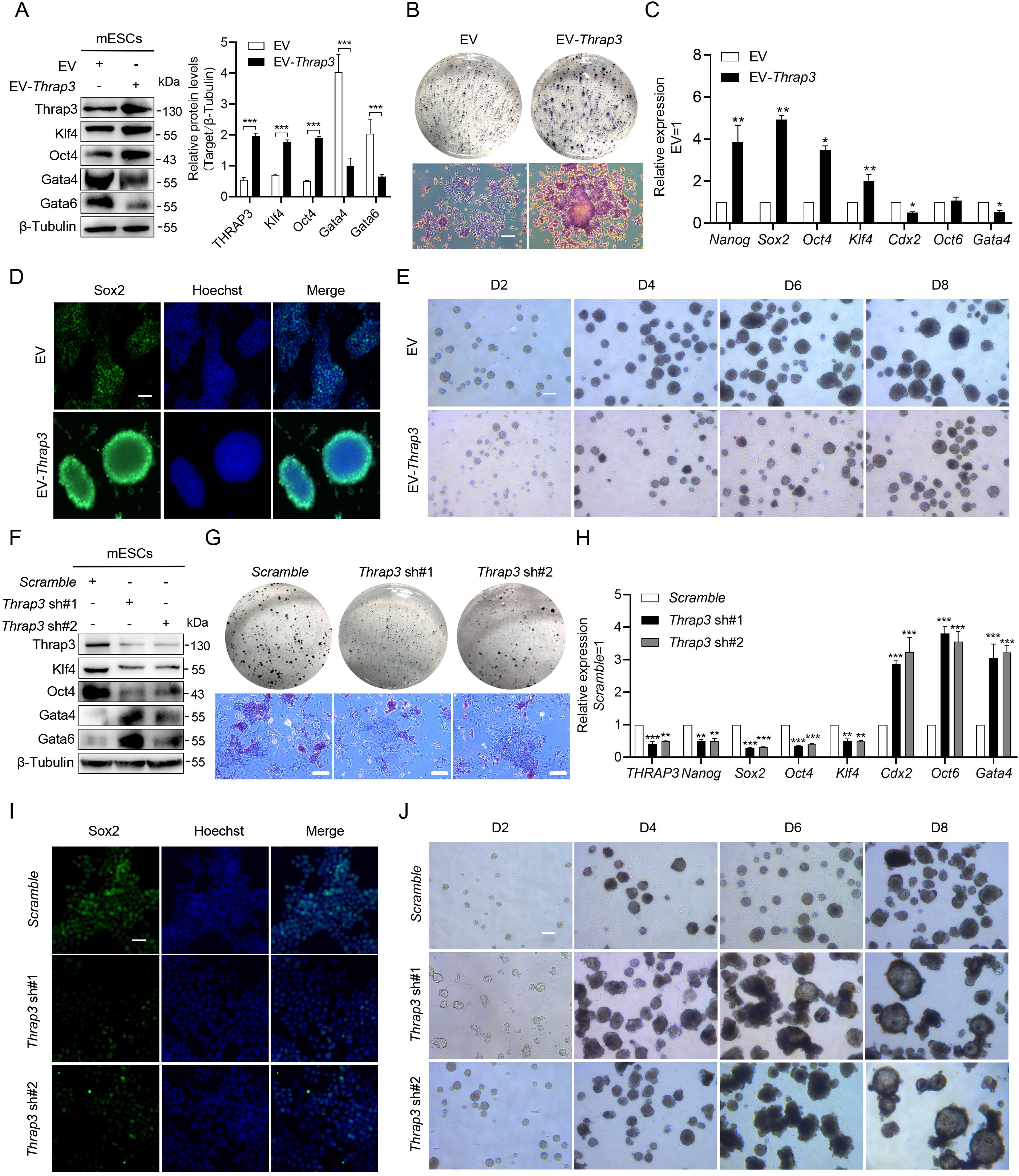
Thrap3 preserves the undifferentiated state in mESCs. **A** Western blot analysis of protein levels of Thrap3, Klf4, Oct4, Gata4 and Gata6 in mESCs transfected with EV or EV-*Thrap3* plasmid after 7 days of culture in the presence of LIF. **B** Colony formation and AP staining of mESCs transfected with EV or *Thrap3*-overexpression plasmid after LIF withdrawal for 7 days (n=3). Scale bar, 50 μm. **C** RT-qPCR analysis of mRNA levels of *Nanog*, *Sox2*, *Oct4*, *Klf4*, *Cdx2*, *Oct6* and *Gata4* in mESCs transfected with EV or *Thrap3*-overexpression plasmid after 7 days of culture in the presence of LIF (n=3). **D** Immunofluorescence staining for Sox2 in mESCs transfected with EV or *Thrap3-*overexpression plasmid after 7 days of culture in the presence of LIF. Scale bar, 50 μm. **E** Representative bright-field images of EB derived from mESCs transfected with EV or *Thrap3*-overexpression plasmids at different days during EB differentiation. Scale bar, 100 μm. **F** Western blot analysis of protein levels of Thrap3, Klf4, Oct4, Gata4 and Gata6 in mESCs transfected with *Scramble* or *Thrap3*-knockdown plasmids after 7 days of culture in the presence of LIF. **G** Colony formation and AP staining assays in mESCs transfected with *Scramble* or *Thrap3*-knockdown plasmids after 7 days of LIF withdrawal (n=3). Scale bar, 50 μm. **H** RT-qPCR analysis of mRNA levels of *Thrap3*, *Nanog*, *Sox2*, *Oct4*, *Klf4*, *Cdx2*, *Oct6* and *Gata4* in mESCs transfected with *Scramble* or *Thrap3*-knockdown plasmids after 7 days of culture in the presence of LIF (n=3). **I** Immunofluorescence staining of Sox2 in mESCs transfected with *Scramble* or *Thrap3*-knockdown plasmids after 7 days of culture in the presence of LIF. Scale bar, 50 μm. **J** Representative images of EB differentiation at different time points derived from mESCs transfected with empty vector *Scramble* or *Thrap3*-knockdown plasmids. Scale bar, 100 μm. The data are presented as mean ± SD. Statistical analyses were performed using two-way ANOVA **A**, **C**, **H**. \**p* < 0.05, \*\**p* < 0.01, \*\*\**p* < 0.001.

We next examined whether *Thrap3* downregulation elicited the opposite phenotype in mESCs maintained in the presence of LIF, using two lentiviral shRNAs (*Thrap3* sh#1 and sh#2). Compared with the *Scramble* control, *Thrap3*-knockdown mESCs exhibited a 60-70% reduction in Thrap3 protein abundance, accompanied by lower Klf4 and Oct4 and higher Gata4 and Gata6 protein levels (Fig. 2F; Fig. S1L). Furthermore, *Thrap3*-knockdown mESCs displayed impaired colony-forming potential and accelerated loss of AP activity following 7 days of LIF withdrawal (Fig. 2G). *Thrap3* knockdown also reduced *Nanog*, *Sox2*, *Oct4* and *Klf4* transcript levels, while increasing *Cdx2*, *Oct6* and *Gata4* transcripts (Fig. 2H), and diminished nuclear Sox2 signal in the presence of LIF (Fig. 2I). Consistently, *Thrap3* knockdown accelerated the loss of *Oct4* and *Sox2* expression and promoted premature differentiation during EB formation (Fig. 2J; Fig. S1M,N). Together, these results support a positive role for Thrap3 in the mESC self-renewal programme.

### 2.3. Thrap3 is required for RNA polymerase II activity on the target genes of Tfcp2l1

To clarify the transcriptional role of Thrap3, we performed RNA sequencing on *Scramble* and *Thrap3* sh#1 mESCs after 7 days of culture in the presence of LIF. Gene Ontology enrichment analysis of differentially expressed genes (DEGs) (|log_2_FC| > 1.0, *p* < 0.05) regulated by *Thrap3* knockdown revealed enrichment of terms related to RNA Pol II transcriptional regulation (Fig. 3A), suggesting that Thrap3 influences RNA Pol II-dependent transcriptional programmes. ChIP-RT-qPCR showed that Thrap3 occupies the same promoter regions of *Esrrb*, *Klf4* and *Prdm14* that are occupied by Tfcp2l1 (Fig. 3B): *Tfcp2l1* overexpression increased Thrap3 enrichment, while Tfcp2l1 depletion reduced enrichment, whereas neither overexpression nor depletion of *Thrap3* altered Tfcp2l1 occupancy (Fig. 3C-F). The dependency is therefore unidirectional. Thrap3 depletion could reduce both Pol II occupancy and Ser2-phosphorylated Pol II CTD across the gene body of the representative target *Esrrb* (Wang *et al*, 2019) (Fig. 3G-J); a Tfcp2l1 N-terminal mutant failed to recruit Thrap3 to the *Esrrb* promoter (Fig. 3K), and *Thrap3* knockdown abolished Tfcp2l1-mediated promoter activation (Fig. 3L). Thrap3 was thus a necessary co-regulatory factor downstream of Tfcp2l1 for efficient Pol II transcription, including establishment of the Ser2P mark.

**Figure 3.**
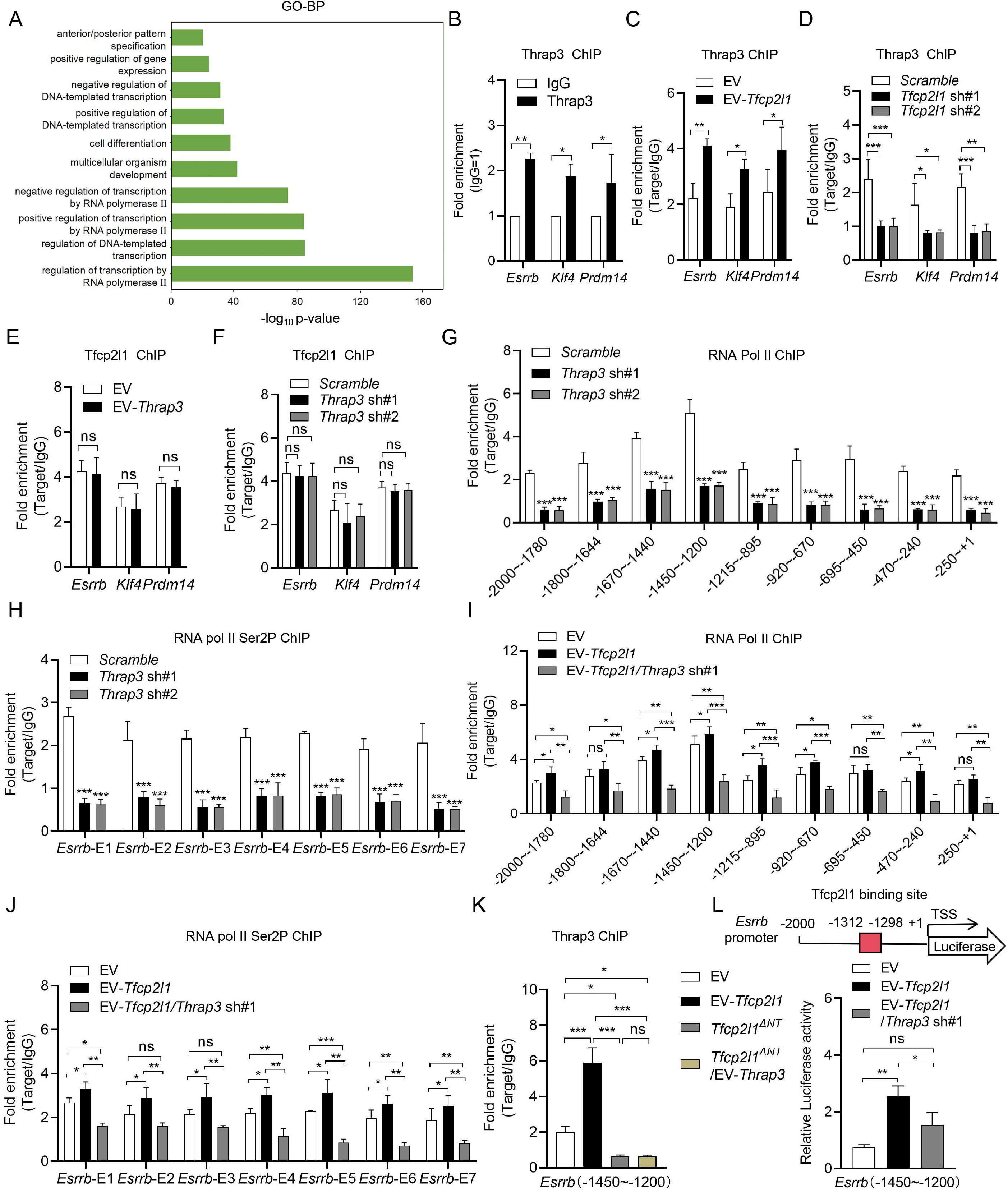
Thrap3 acts as a transcriptional co-regulator to control the downstream targets of Tfcp2l1. **A** mRNA sequencing was performed on mESCs transfected with *Scramble* or *Thrap3* sh#1 plasmid. Differentially expressed genes were screened and subjected to Gene Ontology biological process (GO-BP) enrichment analysis. **B** ChIP-RT-qPCR analysis of Thrap3 enrichment at the designated identical promoter regions of *Esrrb*, *Klf4* and *Prdm14*, key downstream target genes of Tfcp2l1 in mESCs (n=3). **C**, **D** ChIP-RT-qPCR analysis of Thrap3 enrichment at the designated identical promoter regions of Tfcp2l1’s key downstream targets *Esrrb*, *Klf4* and *Prdm14* in mESCs transfected with EV or *Tfcp2l1*-overexpression plasmid (C), and mESCs transfected with *Scramble* or *Tfcp2l1*-knockdown plasmids (D) (n=3). **E**, **F** ChIP-RT-qPCR analysis in mESCs transfected with EV or *Thrap3*-overexpression plasmid (E), as well as *Scramble* or *Thrap3*-knockdown plasmids (F), for the effect on Tfcp2l1 enrichment at the designated promoter regions of its key downstream target genes *Esrrb*, *Klf4* and *Prdm14* (n=3). **G**, **H** ChIP-RT-qPCR analysis of RNA Pol II (G) and Ser2-phosphorylated RNA Pol II CTD (H) enrichment at the *Esrrb* promoter and exons (1-7) in mESCs transfected with *Scramble* or *Thrap3*-knockdown plasmids (n=3). **I**, **J** ChIP-RT-qPCR analysis of the enrichment of RNA Pol II (I) and Ser2-phosphorylated RNA Pol II CTD (J) on the *Esrrb* promoter and exons (1-7) in mESCs transfected with EV, *Tfcp2l1*-overexpression or *Thrap3*-knockdown plasmids (n=3). **K** ChIP-RT-qPCR analysis of Thrap3 enrichment at the *Esrrb* promoter in mESCs transfected with EV, EV-*Tfcp2l1*, *Tfcp2l1^ΔNT^* and EV-*Thrap3* expression plasmids (n=3). **L** Luciferase activity analysis of the *Esrrb* promoter reporter plasmid in mESCs transfected with EV, *Tfcp2l1*-overexpression and *Thrap3*-knockdown plasmids (n=3). Data are presented as mean ± SD. Statistical analyses were performed using two-way ANOVA **B**-**J** and one-way ANOVA **K**, **L**. \**p* < 0.05, \*\**p* < 0.01, \*\*\**p* < 0.001, ns, not significant.

### 2.4. Tfcp2l1 and Thrap3 cooperates to control AS of *Jmjd1c*

RNA Pol II Ser2 phosphorylation is an important marker for the recruitment of the co-transcriptional splicing machinery (Egloff & Murphy, 2008; Jonkers & Lis, 2015). To examine the effects of *Tfcp2l1* or *Thrap3* knockdown on AS, we performed rMATS analysis on RNA-seq data from *Tfcp2l1*- or *Thrap3*-knockdown mESCs, covering five classes of splicing events: exon skipping (SE), mutually exclusive exons (MXE), alternative 5′ splice sites (A5SS), alternative 3′ splice sites (A3SS) and intron retention (RI). Using FDR < 0.05 and |ΔPSI| > 0.2 as thresholds, we identified 276 shared genes and 624 shared events; 88.94% (555/624) changed in the same direction after either knockdown (Fig. 4A; Fig. S2A). Among them, six genes with multiple AS patterns (*Jmjd1c*, *Eif4a2*, *Tia1*, *Ccdc14*, *Mef2* and *Ncf2*) have been implicated in ESCs (Fig. 4B; Supplementary Table 2). We selected Jmjd1c for detailed analysis because of its reported roles in stem-cell maintenance and the DNA-damage response, whereas its splicing regulation has not been well characterized in ESCs.

**Figure 4.**
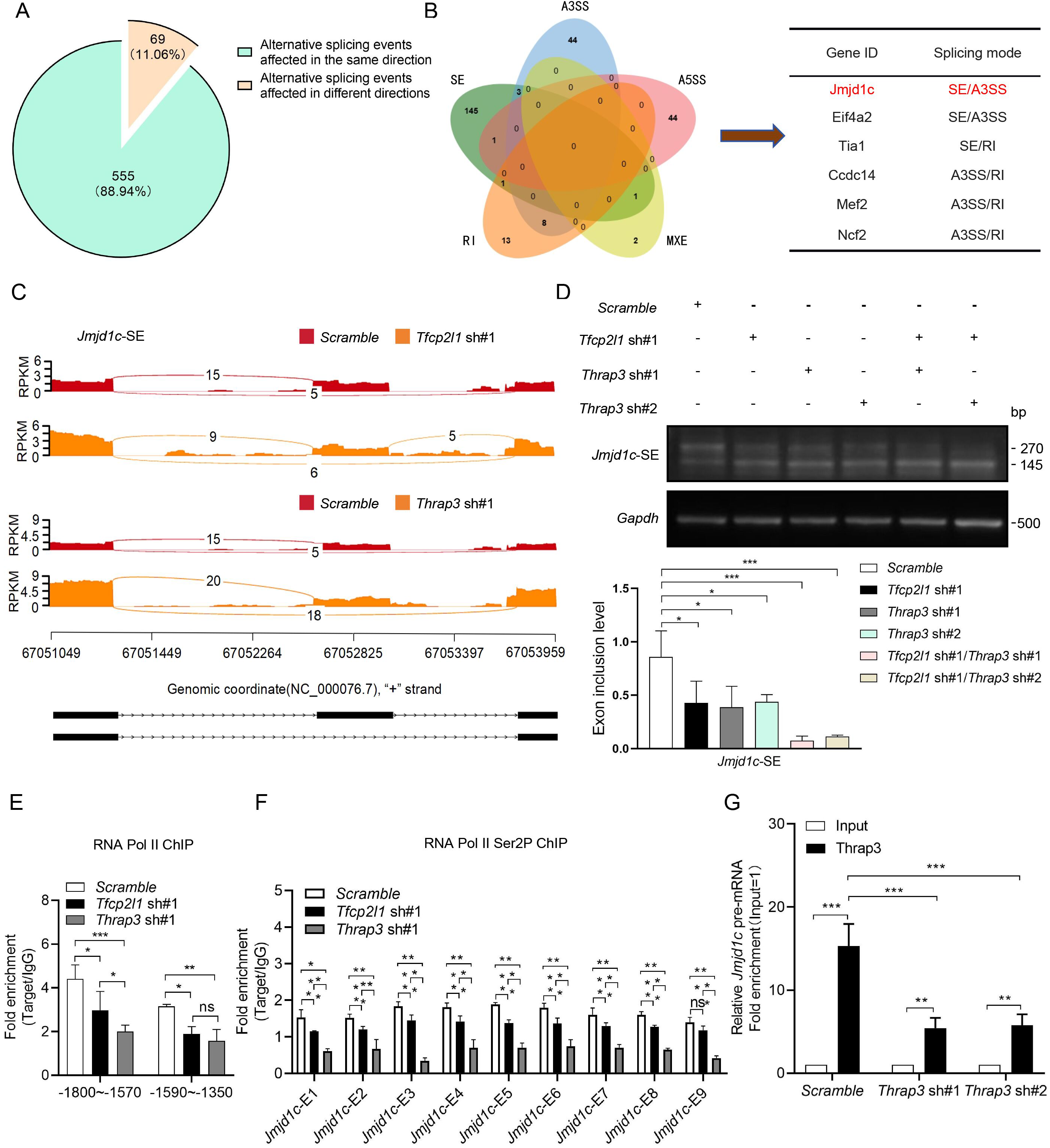
Knocking down *Tfcp2l1* or *Thrap3* promotes *Jmjd1c* SE. **A** Pie chart illustrating alternative splicing events regulated by Tfcp2l1 and Thrap3 in mESCs. **B** Overlap of five types of alternative splicing events in *Tfcp2l1*-knockdown or *Thrap3*-knockdown mESCs, compared with *scramble* control. **C** Representative Sashimi plots of *Jmjd1c* exon skipping (SE) events analyzed by rMATS in cells transfected with *Scramble*, *Tfcp2l1* sh#1 or *Thrap3* sh#1. **D** RT-PCR analysis of Jmjd1c exon skipping in *Scramble*, *Tfcp2l1*-knockdown or *Thrap3*-knockdown mESCs. The histogram shows quantitative results of RT-PCR assays (n=3). **E** ChIP-RT-qPCR analysis of RNA Pol II enrichment at the *Jmjd1c* promoter in *Scramble*, *Tfcp2l1*-knockdown or *Thrap3*-knockdown mESCs (n=3). **F** ChIP-RT-qPCR analysis of the enrichment of Ser2-phosphorylated RNA Pol II CTD at the Jmjd1c exons (1-9) region in *Scramble*, *Tfcp2l1*-knockdown or *Thrap3*-knockdown mESCs (n=3). **G** RIP-RT-qPCR analysis of Thrap3 enrichment on pre-mRNA spanning the exon-5 region of Jmjd1c in *Scramble* or *Thrap3*-knockdown mESCs (n=3). Data are presented as mean ± SD. Statistical analyses were performed using one-way ANOVA **D** and two-way ANOVA **E**, **F**, **G**. \**p* < 0.05, \*\**p* < 0.01, \*\*\**p* < 0.001, ns, not significant.

Sashimi plots and isoform-specific reverse-transcription PCR (RT-PCR) confirmed prominent skipping of *Jmjd1c* exon-5 upon *Tfcp2l1* or *Thrap3* knockdown, accompanied by altered A3SS usage and reduced percent-spliced-in (PSI) values (Fig. 4C,D; Fig. S2B,C). To investigate whether Tfcp2l1 and Thrap3 bind to the Jmjd1c locus and regulate its AS, we performed ChIP-RT-qPCR, which revealed that Tfcp2l1 and Thrap3 were enriched in the same region of the Jmjd1c promoter (Fig. S2D,E). *Tfcp2l1* knockdown reduced Thrap3 enrichment, whereas *Thrap3* knockdown did not alter Tfcp2l1 enrichment (Fig. S2F,G). *Tfcp2l1* overexpression increased *Jmjd1c* promoter-reporter activity, and this effect depended on Thrap3 (Fig. S2H). Knockdown of either factor reduced RNA Pol II enrichment at the *Jmjd1c* promoter and the RNA Pol II Ser2P signal across the gene body (Fig. 4E,F). RIP-RT-qPCR suggested that Tfcp2l1 and Thrap3 associate with *Jmjd1c* pre-mRNA containing the exon-5 region (Fig. 4G; Fig. S2I). Total *Jmjd1c* mRNA abundance also decreased after *Tfcp2l1* or *Thrap3* knockdown (Fig. S2J,K). These results suggest that Tfcp2l1 and Thrap3 influence both total *Jmjd1c* transcript abundance and exon-5 inclusion.

### 2.5. Loss of Tfcp2l1 or Thrap3 triggers DNA damage accumulation and apoptosis

To explore whether *Tfcp2l1* or *Thrap3* knockdown genuinely induces DNA damage and accelerates apoptosis, we first assessed γ-H2AX (Prabhu *et al*, 2024) foci by immunofluorescence. Both knockdown groups revealed markedly higher numbers of nuclear γ-H2AX foci (Fig. 5A,B), whereas Atm and Atr mRNA levels and the total protein abundance of Bclaf1, Atm, p-Atm^S1981^, Atr and p-Atr^S428^ were largely unchanged (Fig. 5C,D; Fig. S3A-D), as was the abundance of Bclaf1, the Thrap3 paralogue, indicating that the phenotype reflects loss of Thrap3 itself rather than a secondary change in the related factor. These findings suggest that the elevated γ-H2AX foci are not explained by changes in the measured expression levels of these DNA damage response kinases.

**Figure 5.**
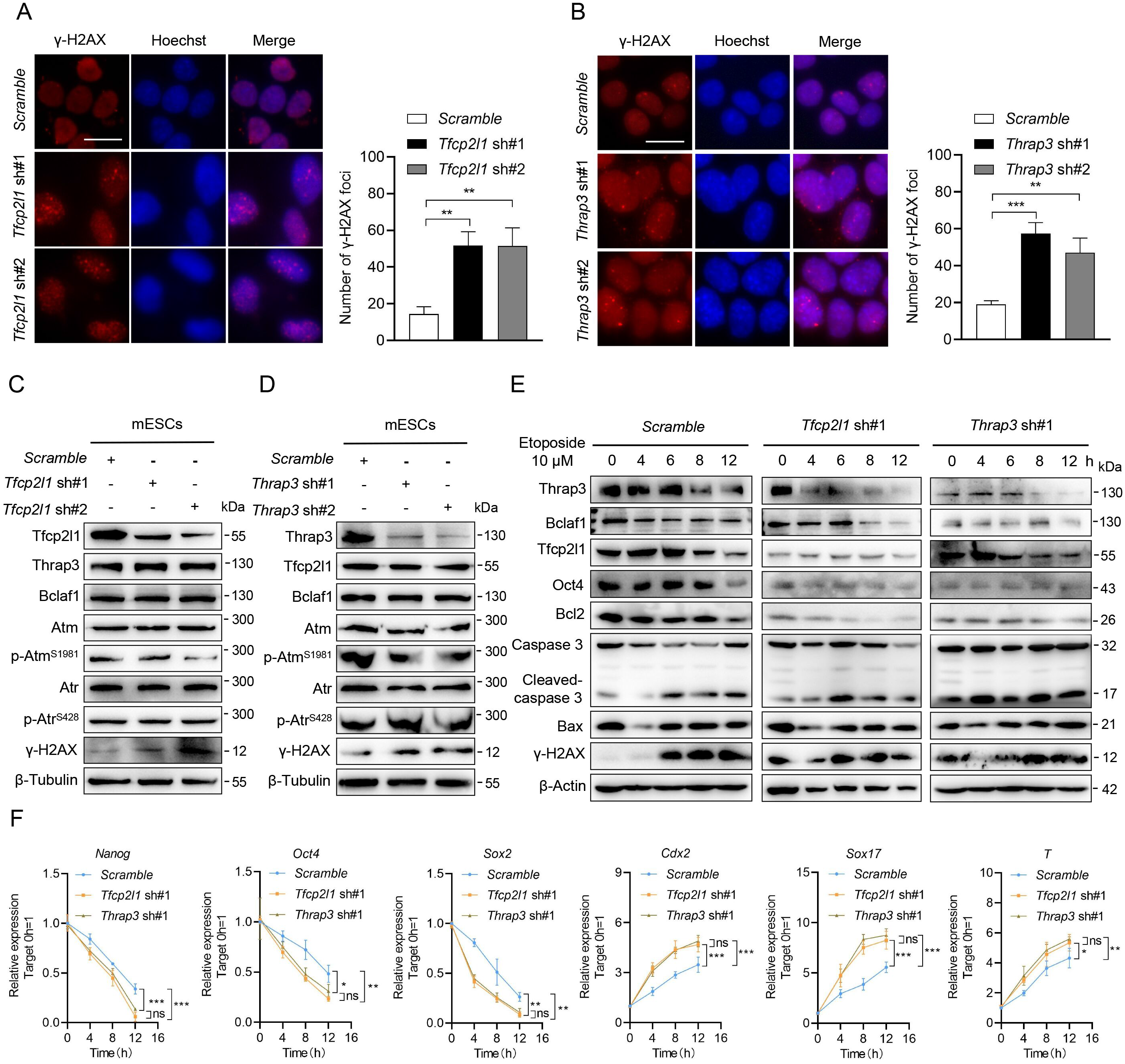
Knockdown of *Tfcp2l1* or *Thrap3* induces DNA damage. **A**, **B** Representative immunofluorescence images of the DNA damage marker γ-H2AX in *Tfcp2l1*-knockdown (A) or *Thrap3*-knockdown (B) mESCs compared with the *Scramble* group. Scale bar, 20 μm. **C**, **D** Western blot analysis of protein levels of Tfcp2l1, Thrap3, Bclaf1, Atm, p-Atm^S1981^, Atr, p-Atr^S428^, and γ-H2AX in mESCs transfected with *Scramble*, *Tfcp2l1*-knockdown (C) or *Thrap3*-knockdown (D) plasmids. **E** Western blot analysis of protein levels of Tfcp2l1, Thrap3, Bclaf1, Oct4, Bcl2, Caspase 3, Cleaved caspase 3, Bax and γ-H2AX in mESCs transfected with *Scramble, Tfcp2l1*-knockdown or *Thrap3*-knockdown plasmids, after treatment with 10 μM etoposide for 0, 4, 6, 8, 12 h. **F** RT-qPCR analysis of mRNA levels of *Nanog*, *Oct4*, *Sox2*, *Cdx2*, *Sox17* and *T* in mESCs. Cells were transfected with *Scramble*, *Tfcp2l1*-knockdown or *Thrap3*-knockdown plasmids and treated with 10 μM etoposide for the indicated time points (n=3). Data are presented as mean ± SD. Statistical analyses were performed using one-way ANOVA (**A**, **B)**, and two-way ANOVA **F**. \**p* < 0.05, \*\**p* < 0.01, \*\*\**p* < 0.001; ns, not significant.

Next, to assess the physiological consequences of DNA damage, we treated cells with 1 μM etoposide, a DNA double-strand-break inducer, and subsequently withdrew LIF. Both knockdown groups showed severely impaired colony formation and AP activity (Fig. S3E). We titrated etoposide to identify a concentration that produced robust acute DNA damage. Addition of 10 μM etoposide induced clear morphological changes and reduced cell viability, whereas lower doses produced weaker effects; we therefore used 10 μM for the time-course experiments (Fig. S3F,G). Under this treatment, Western blot results showed that *Tfcp2l1* or *Thrap3* knockdown led to reduced Oct4 and Bcl2 protein levels, increased Bax levels, and accumulation of Cleaved caspase 3 and γ-H2AX (Fig. 5E), whereas differentiation-marker expression increased (Fig. 5F). These results suggest that Tfcp2l1 or Thrap3 deficiency increases susceptibility to genotoxic stress and is associated with the loss of self-renewal markers and apoptosis-associated changes.

### 2.6. The exon-5 of *Jmjd1c* is essential for the interaction between Mdc1 and Rnf8

Previous work has shown that human JMJD1C promotes MDC1 and RNF8 binding by demethylating MDC1 at K45 (Watanabe *et al*, 2013). Since knockdown of *Tfcp2l1* or *Thrap3* suppresses inclusion of the exon-5 coding region, we hypothesized that this region is essential for the interaction of Jmjd1c and Mdc1. Endogenous co-immunoprecipitation confirmed Jmjd1c and Mdc1 association in mESCs (Fig. 6A), and a panel of truncation constructs showed that deleting the exon-5 coding region (Jmjd1c-ΔE5) weakened Mdc1 binding (Fig. 6B,C). Immunoprecipitation with a Pan Dimethyl-lysine antibody showed that Mdc1 dimethylation was increased in cells expressing *Jmjd1c*-ΔE5; this effect was detected under control conditions and persisted upon exposure to Methylstat, a JmjC-domain demethylase inhibitor (Fig. 6D). Overexpression of *Jmjd1c*-ΔE5 also reduced Rnf8 and K63-linked ubiquitin chains co-precipitated with Mdc1 (Fig. 6E). Consistent with this finding, knockdown of *Tfcp2l1* or *Thrap3* recapitulated this phenotype without altering total Mdc1 or Rnf8 mRNA and protein levels (Fig. S4A,B), localising the defect to protein modification and interaction rather than protein abundance. Sequence alignment of human and mouse Mdc1 identified mouse Mdc1 K53 as the orthologous residue corresponding to the conserved human K45 site (Supplementary Table 3). The *Mdc1^K53A^* mutant enhanced Mdc1 and Rnf8 association, and 10 μM etoposide treatment did not further alter this interaction (Fig. 6F), indicating that modification of this residue restrains the Mdc1 and Rnf8 interaction and is not a downstream consequence of DNA damage. Immunofluorescence confirmed that knockdown of *Tfcp2l1* or *Thrap3* reduced Mdc1 and Rnf8 co-localization at DNA damage sites after 4 h of treatment with 10 μM etoposide (Fig. S4C). Similarly, *Jmjd1c*-ΔE5 expression or Methylstat treatment diminished this co-localization (Fig. S4D). In summary, these data identify the Jmjd1c exon-5 region as a key determinant governing Mdc1 demethylation and place this event upstream of the Mdc1 and Rnf8-dependent K63-linked ubiquitination signal.

**Figure 6.**
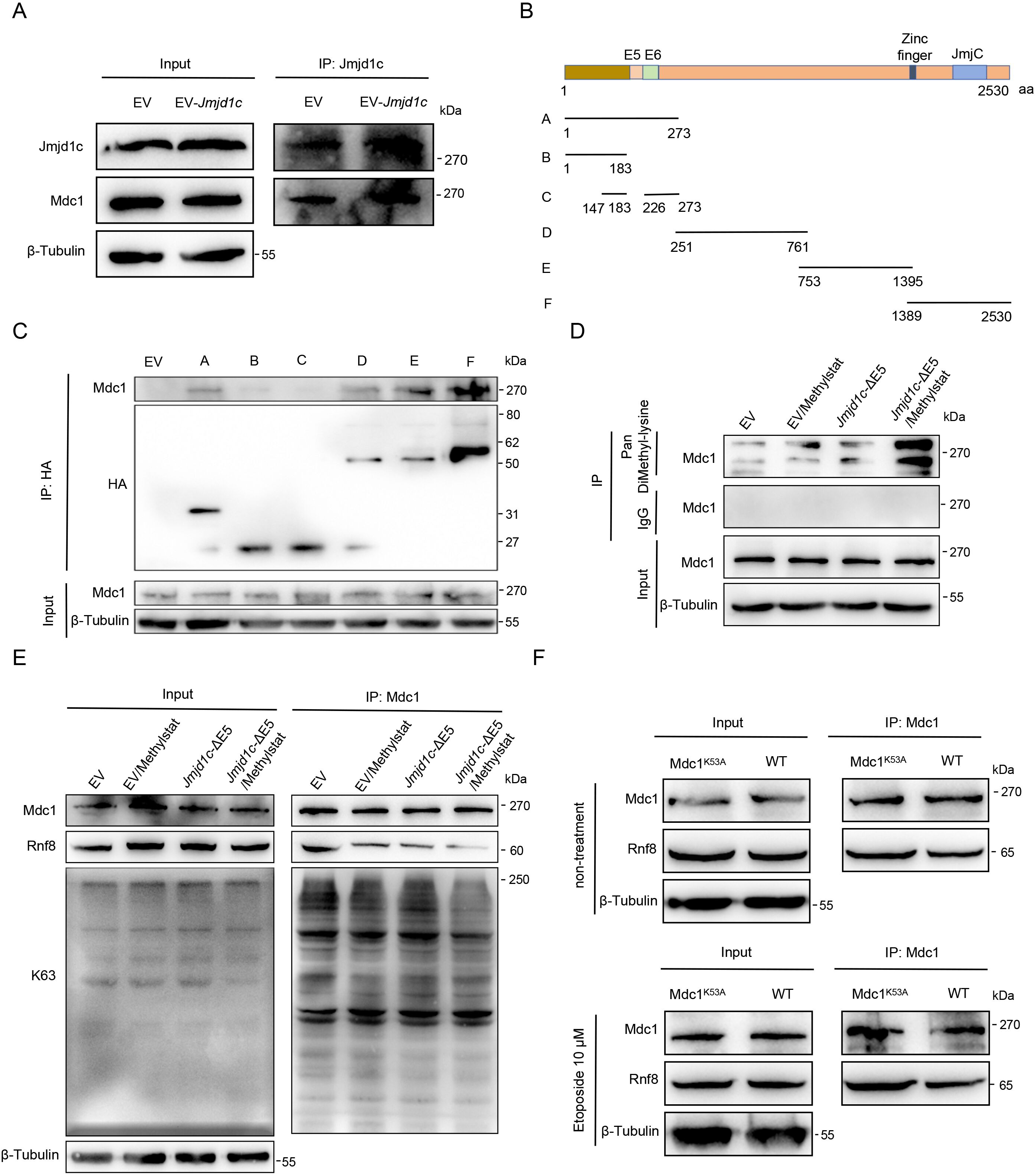
*Jmjd1c*-E5 deletion attenuates the binding of Jmjd1c, Mdc1 and Rnf8 via impairing Mdc1 demethylation. **A** Co-IP assay detecting the interaction between Jmjd1c and Mdc1 in mESCs. **B** Schematic diagram of plasmid constructs expressing truncated Jmjd1c protein variants. **C** Co-IP assay examining the interaction of Mdc1 with different truncated Jmjd1c variants after transfection of corresponding plasmids in mESCs. **D** Immunoprecipitation with Pan Dimethyl-lysine antibody in mESCs transfected with empty vector (EV) or the *Jmjd1c*-ΔE5 construct, with or without Methylstat (5 μM) treatment. **E** Co-IP assay with Mdc1-antibody to detect its binding to Rnf8 and K63-linked ubiquitin chains in mESCs transfected with EV or *Jmjd1c*-ΔE5, in the presence or absence of the JmjC domain inhibitor Methylstat (5 μM). **F** Co-IP assay detecting the interaction between Mdc1 and Rnf8 in wild-type (WT) mESCs and mESCs transfected with *Mdc1^K53A^* plasmid, with or without 10 μM etoposide treatment for 4 h.

### 2.7. Loss of Tfcp2l1 or Thrap3 inhibits Rap80 and Brca1 recruitment to DNA damage sites and increases Rad51 foci formation

To examine downstream DNA-damage signalling, we analysed total protein extracts from *Tfcp2l1*- or *Thrap3*-knockdown mESCs. Knockdown of either Tfcp2l1 or Thrap3 led to reduced Jmjd1c protein abundance, whereas the protein levels of Rap80 and Brca1, core components modulating HR-dependent repair (Kim *et al*, 2007; Zhang *et al*, 2012), remained unchanged. Meanwhile, 53bp1, a key regulatory protein that favours NHEJ-mediated repair, was only mildly affected (Chapman *et al*, 2013; Makins *et al*, 2025) (Fig. 7A,B; Fig. S4E,F). The differences were more pronounced at damaged chromatin. After 4 h of treatment with 10 μM etoposide, numbers of Brca1 foci were reduced in both knockdown backgrounds, whereas 53bp1 recruitment remained unaffected (Fig. 7C; Fig. S4G). In *Tfcp2l1*- or *Thrap3*-knockdown mESCs, levels of chromatin-bound Jmjd1c, Rnf8, Rap80, Brca1 protein and K63-linked ubiquitin chains decreased upon 10 μM etoposide treatment for 0, 4 and 8 h, whereas chromatin-bound 53BP1 levels remained largely unchanged (Fig. 7D). Numbers of Rad51 foci increased after 4 h of 10 μM etoposide treatment in both knockdown backgrounds (Fig. 7E; Fig. S4H). These findings indicate that aberrant *Jmjd1c* splicing upon *Tfcp2l1* or *Thrap3* knockdown perturbs the Rap80-Brca1 arm of the DNA damage response, with no obvious impact on the 53bp1-dependent branch. This leads to dysregulated DDR signalling and persistent DNA damage accumulation.

**Figure 7.**
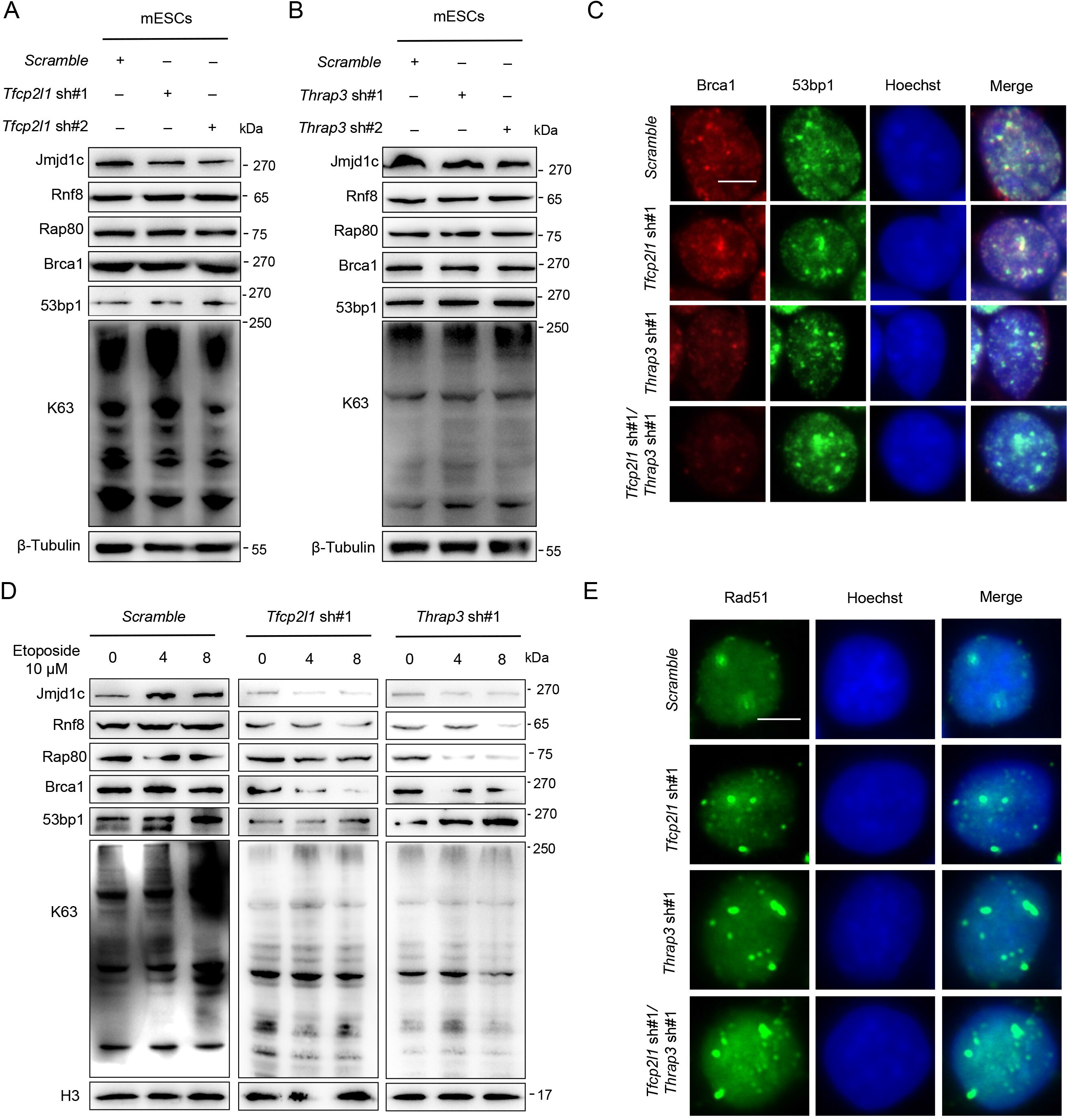
Knockdown of *Tfcp2l1* or *Thrap3* impairs the accumulation of Rap80 and Brca1 on chromatin following DNA damage. **A**, **B** Western blot analysis of protein levels of Jmjd1c, Rnf8, Rap80, Brca1, 53bp1 and K63-linked ubiquitin chains in mESCs transfected with *Scramble*, *Tfcp2l1*-knockdown (A) or *Thrap3*-knockdown (B) plasmids. **C** Representative immunofluorescence images of nuclear Brca1 (red) and 53bp1 (green) in mESCs transfected with *Scramble*, *Tfcp2l1*-knockdown or *Thrap3*-knockdown plasmids, after 4 h treatment with 10 μM etoposide. Scale bar, 10 μm. **D** Western blot analysis of Jmjd1c, Rnf8, Rap80, Brca1, 53bp1 and K63-linked ubiquitin chains in chromatin-bound protein extracts from mESCs. Cells in *Scramble*, *Tfcp2l1*-knockdown or *Thrap3*-knockdown groups were treated with 10 μM etoposide for 0, 4 and 8 h, respectively. **E** Representative immunofluorescence images of nuclear Rad51 in mESCs transfected with *Scramble*, *Tfcp2l1*-knockdown or *Thrap3*-knockdown plasmids, after 4 h treatment with 10 μM etoposide. Scale bar, 10 μm.

### 2.8. Conserved TFCP2L1- and THRAP3-mediated regulation of DDR repair pathway choice in hESCs

To verify the conservation of these key findings in hESCs, we performed endogenous Co-IP, which confirmed the TFCP2L1 and THRAP3 interaction in hESCs; immunofluorescence further showed their nuclear co-localization. The same critical residues are required for binding: residues F3, P8, and P27 in TFCP2L1 and R401, K410, and R749 in THRAP3 are essential for their association in hESCs (Fig. S5A-F). *THRAP3* overexpression elevated KLF4 and OCT4 protein levels, reduced GATA4 and GATA6 protein levels, enhanced colony-formation capacity and AP activity, and increased self-renewal gene expression as well as SOX2 immunoreactivity (Fig. S5G-K), whereas THRAP3 depletion reduced hESC self-renewal (Fig. S6A-D). These data confirm a direct interaction between TFCP2L1 and THRAP3 in hESCs, and THRAP3 also supports hESC self-renewal.

At the splicing level, depletion of TFCP2L1 or THRAP3 promoted SE and A3SS AS events of *JMJD1C* (Fig. S7A,B); both factors occupy the same promoter region and exhibit unidirectional dependency (Fig. S7C-H); and both bind exon-5-spanning pre-mRNA (Fig. S7I,J). Total *JMJD1C* mRNA abundance and promoter reporter activity decreased upon knockdown of either factor (Fig. S7K-M). These results indicate that TFCP2L1 and THRAP3 regulate AS of *JMJD1C* in hESCs. Moreover, depletion of either factor increased numbers of γ-H2AX foci (Fig. 8A), elevated MDC1 dimethylation (Fig. 8B), and reduced RNF8 and K63-linked ubiquitin chains bound to MDC1 without altering total MDC1 or RNF8 protein levels (Fig. 8C; Fig. S8A-D). After treatment with 10 μM etoposide, self-renewal markers were downregulated, and apoptotic markers accumulated (Fig. S8E). In addition, etoposide did not alter the regulatory role of MDC1 K45 in the MDC1 and RNF8 interaction (Fig. 8D). Meanwhile, numbers of BRCA1 and RAP80 foci decreased, whereas 53BP1 foci remained unchanged and RAD51 foci increased (Fig. 8E,F; Fig. S8F-I). These results demonstrate conservation of this regulatory cascade controlled by TFCP2L1 and THRAP3 across mouse and human ESCs.

**Figure 8.**
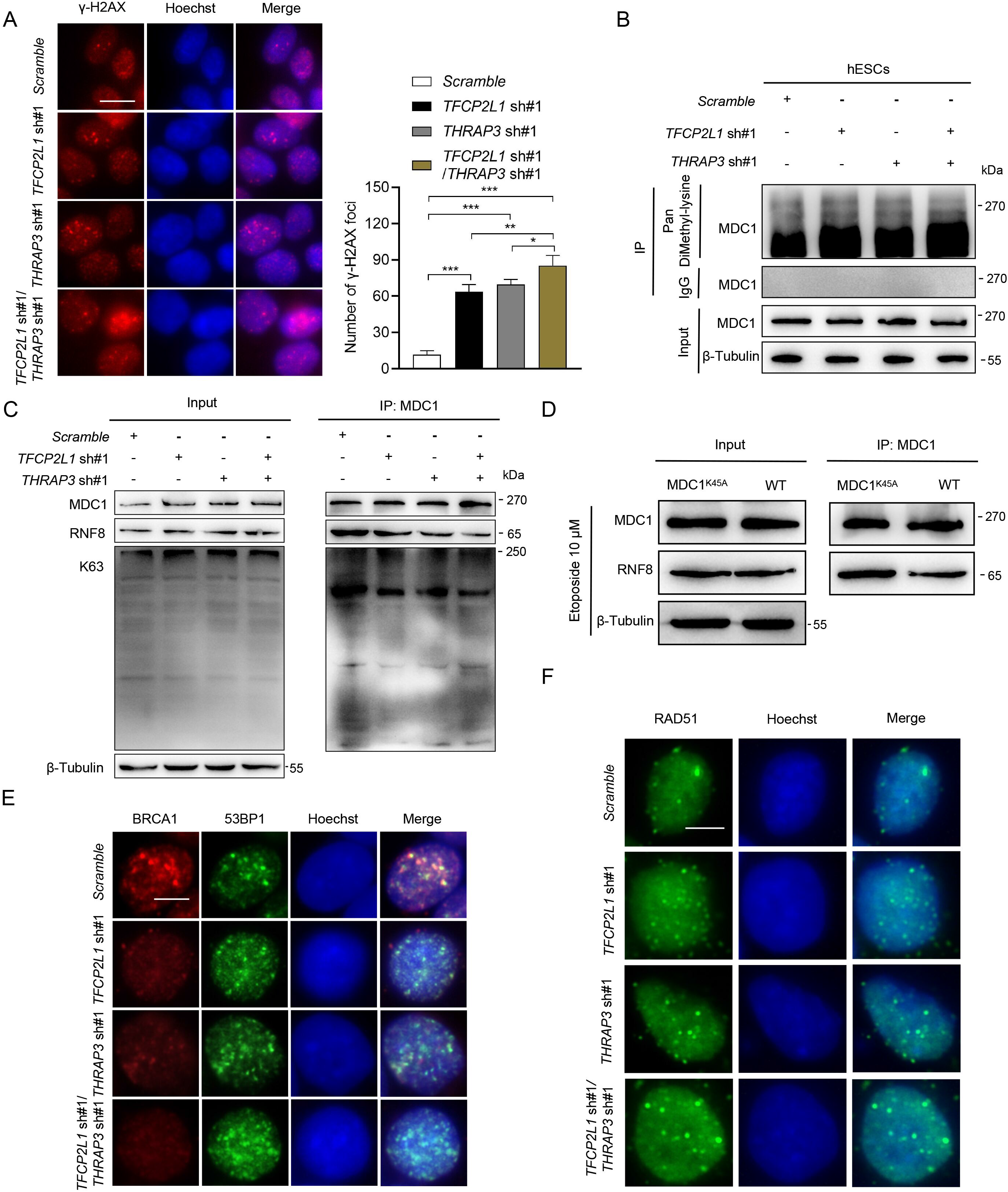
Knockdown of *TFCP2L1* or *THRAP3* in hESCs inhibits MDC1 demethylation and hinders BRCA1 recruitment to damage sites. **A** Representative immunofluorescence images of the DNA damage marker γ-H2AX in hESCs transfected with *Scramble*, *TFCP2L1*-knockdown or *THRAP3*-knockdown plasmids. Scale bar, 20 μm. **B** Immunoprecipitation with Pan DiMethyl-lysine antibody in hESCs transfected with *Scramble*, *TFCP2L1*-knockdown or *THRAP3*-knockdown plasmids. **C** Co-IP assay using MDC1-antibody to detect the binding of MDC1 to RNF8 and K63-linked ubiquitin chains in hESCs transfected with *Scramble*, *TFCP2L1*-knockdown or *THRAP3*-knockdown plasmids. **D** Co-IP assay detecting the interaction between MDC1 and RNF8 in wild-type (WT) hESCs and hESCs transfected with *MDC1^K45A^* plasmid after treatment with 10 μM etoposide for 4 h. **E** Representative immunofluorescence images of nuclear BRCA1 (red) and 53BP1 (green) in hESCs transfected with *Scramble*, *TFCP2L1*-knockdown or *THRAP3*-knockdown plasmids, following 4 h treatment with 10 μM etoposide. Scale bar, 10 μm. **F** Representative immunofluorescence images of nuclear RAD51 in hESCs transfected with *Scramble*, *TFCP2L1*-knockdown or *THRAP3*-knockdown plasmids, following 4 h treatment with 10 μM etoposide. Scale bar, 10 μm. Data are presented as mean ± SD. Statistical analyses were performed using one-way ANOVA **A** . \**p* < 0.05, \*\**p* < 0.01, \*\*\**p* < 0.001.

## 3. Discussion

Tfcp2l1 has been studied almost entirely as a DNA-binding transcriptional regulator. Our affinity purification identified thyroid hormone receptor associated protein 3 (Thrap3), a cofactor of the transcriptional elongation machinery (Vohhodina *et al*, 2017; Yarosh *et al*, 2015). The same N-terminal segment, which we identified here as the Thrap3 docking site, is also required for self-renewal and mediates Mta1 recruitment, indicating that it is a general docking surface for co-regulators rather than a Tfcp2l1-specific motif. Because the docking element lies outside the CP2 DNA-binding domain (Kim *et al*, 2016), Thrap3 association does not displace Tfcp2l1 from chromatin; Tfcp2l1 instead operates as a platform that delivers splicing machinery to its own target loci. The dependency between the two proteins is strictly unidirectional: Thrap3 recruitment to target promoters depends on Tfcp2l1, whereas Tfcp2l1 occupancy is unaffected by Thrap3 depletion. This places Tfcp2l1 upstream of both elongation and co-transcriptional splicing and explains why loss of Thrap3 compromises Ser2P-dependent transcription at pluripotency loci without a reciprocal effect on Tfcp2l1 binding. We expand the self-renewal network beyond promoter-directed transcription to a mechanism that co-regulates RNA processing and, in turn, governs DNA repair pathway choice in ESCs (Fig. 9).

**Figure 9.**
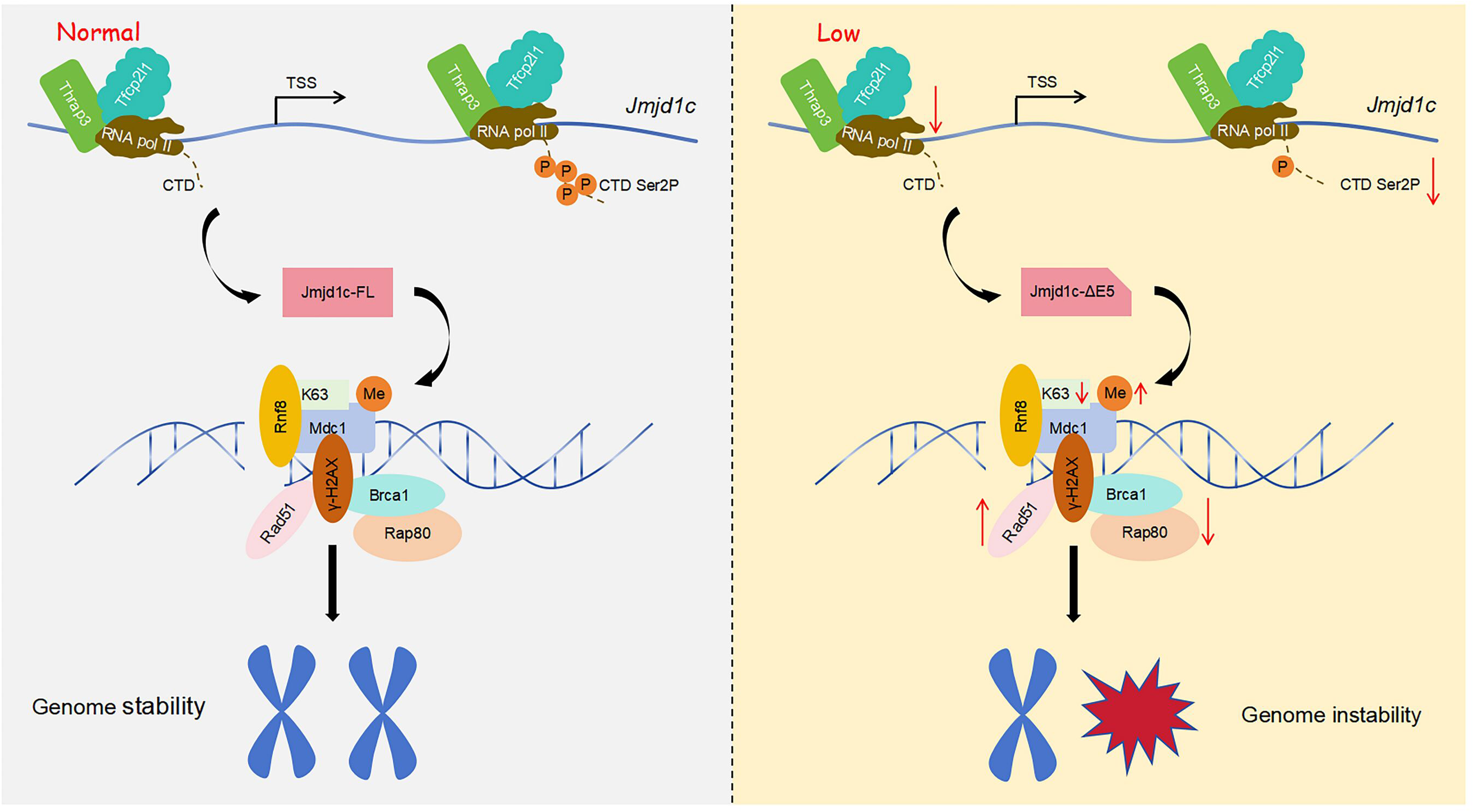
Schematic diagram showing the complex of Tfcp2l1 and Thrap3 in maintaining ESC genomic homeostasis. Loss of function of Tfcp2l1 and Thrap3 in ESCs induces *Jmjd1c* exon-5-skipping, disrupts the Mdc1-Rnf8-Rap80-Brca1 signaling cascade, and leads to dysregulated DNA damage response and impaired genomic homeostasis. This regulatory axis is conserved in both mouse and human ESCs.

THRAP3 and its paralog BCLAF1 regulate AS and nuclear export of transcripts encoding DNA repair proteins; their depletion confers sensitivity to DNA-damaging agents, impaired DNA repair and genomic instability (Vohhodina *et al*., 2017). This function operates independently of DNA damage, as a distinct set of transcripts is processed constitutively. Our findings do not establish that Thrap3 is recruited in response to DNA damage; instead, we identify Tfcp2l1 as the factor responsible for recruiting Thrap3 in undifferentiated ESCs. Genome-scale screens likewise identified RBMX as a splicing-dependent component of the HR response (Adamson *et al*, 2012), indicating that RNA processing is a general, rather than incidental, layer of pathway regulation. Our data place Thrap3 upstream of these functional outputs, rather than placing it in a parallel pathway. Depletion of either factor alters 624 shared splicing events, 88.94% of which change in the same direction, indicating that the two proteins function as a single regulatory unit rather than parallel regulators. The repair programme is therefore specified by the pluripotency circuitry while cells remain undifferentiated and damage-free; AS should in this sense be regarded as a routine output of stemness circuitry rather than a specialized response to damage. Co-transcriptional factors such as Thrap3 belong to the pluripotency apparatus rather than the auxiliary transcription factor toolbox. Pre-establishing such a programme is an economical strategy for ESCs, whose short G1 phase and weak checkpoints permit continuous generation of replication-associated lesions (Kurashima *et al*., 2024; Liu *et al*., 2019). This model also reverses conventional causality, in that the transcriptional programme is not merely a downstream beneficiary of repair status but an upstream determinant of it.

Jmjd1c serves as the convergence point for both arms of this circuit. Beyond its roles in stem cell transcription and somatic reprogramming, JMJD1C demethylates MDC1 at K45 (K53 in mouse) (Watanabe *et al*., 2013; Xiao *et al*, 2017), which relieves inhibition of RNF8 docking and enables assembly of the RAP80 and BRCA1 repair branch. By contrast, 53BP1 and RIF1 redirect this same substrate toward classical NHEJ (Chapman *et al*., 2013; Makins *et al*., 2025). Pathway choice is therefore determined at the level of an early chromatin modification, not by the repair machineries themselves. Our data further show that this decision is controlled by transcript maturation: the exon-5-containing isoform is required for Jmjd1c to interact with Mdc1; its absence leaves the ubiquitin cascade entirely unassembled rather than merely attenuated. Alanine substitution at K53 strengthened Mdc1 and Rnf8 association and rendered it refractory to etoposide, establishing that this residue controls their binding. Methyl-mimic constructs will be required to determine whether methylation at this site is inhibitory, as Pan Dimethyl-lysine antibodies cannot assign the signal to a residue. This shifts repair pathway bias rather than enhancing homologous recombination, as might be inferred from increased Rad51 foci. Reduced Rap80 and Brca1 occupancy removes the constraint Brca1 imposes on end resection (Feng *et al*, 2026), allowing resected intermediates to accumulate and recruit Rad51. Consistent with published observations of JMJD1C loss, BRCA1 and BARD1 then protect replication forks rather than promoting their degradation (Daza-Martin *et al*, 2019; Densham *et al*, 2016). The unchanged abundance and phosphorylation of ATM and ATR place this defect downstream of damage sensing. This module extends beyond DNA repair biochemistry to govern life-or-death cell fate decisions. Undifferentiated ESCs maintain high apoptotic priming through constitutive expression of the BH3-only protein NOXA, which antagonizes MCL1 to eliminate irreparably damaged cells (Basundra *et al*, 2022). The module therefore sets both repair fidelity and the threshold for damage-induced apoptosis, so that when repair capacity is compromised, cells that would normally be cleared survive and expand (Szmyd *et al*, 2025; Weissbein *et al*., 2014; Zhang *et al*, 2019), which explains how culture-acquired genomic abnormalities accumulate and propagate in ESC populations prone to recurrent chromosomal gains during prolonged in vitro culture.

In summary, we uncovered a Tfcp2l1 function independent of its canonical DNA-binding transcriptional activity and defined the structural basis underlying this function. We describe an RNA-processing mechanism through which this function is exerted, linking AS of a single transcript to the recruitment of DNA-repair-associated factors, and show that key features of this circuit are conserved in human embryonic stem cells. Several open questions remain, including which trans-acting factors translate RNA Pol II Ser2P status into *Jmjd1c* exon-5 splicing decisions and whether this regulatory circuit operates under physiological endogenous replication stress. Answering these questions will advance our understanding of how genomic fidelity is maintained during pluripotent stem cell expansion.

## 4. Materials and methods

### Cell lines and cell culture

The mESCs and human embryonic kidney HEK-293T cell line used in this study were purchased from the Cell Bank of the Chinese Academy of Sciences (Shanghai, China). Cells were seeded onto 0.1% gelatin-coated 6-well plates (Cat. 11110, Labselect, China). Cells were cultured in Dulbecco’s Modified Eagle Medium (DMEM, Transgene Biotech, China) supplemented with 10% fetal bovine serum (FSP500; ExCell Bio, Australia), 1% penicillin-streptomycin, 1% non-essential amino acids (11140-050, GIBCO, USA), and 1% β-mercaptoethanol (M3148, Sigma). In addition, 1000 U/mL LIF (LIF1010, Millipore, USA) was supplemented to maintain pluripotency. Cells were incubated in a humidified incubator at 37 °C with 5% CO₂. The presence of mycoplasma was tested in all cell lines every month to rule out mycoplasma infection.

### Plasmid construction

The coding sequences of *Tfcp2l1*, *Thrap3*, *Jmjd1c* and *Mdc1* were inserted into the PiggyBac transposon vector bearing FLAG and HA tags. Mutant plasmids were constructed by overlap-extension PCR. For shRNA plasmids, the pLKO.1-TRC vector (10878, Addgene) was used. Specific shRNA sequences targeting *Tfcp2l1* and *Thrap3* were inserted between the AgeI and EcoRI restriction sites. Primer sequences and shRNA sequences are listed in Table S1 and Table S2.

### Cell transfection and lentiviral infection

For lentivirus packaging, 2 μg of target gene plasmid, together with packaging plasmids psPAX2 (1.25 μg) and pMD2.G (0.75 μg), were co-transfected into HEK-293T cells using 8 μg of liposomal transfection reagent (40802ES03, Yeasen Biotechnology, China). Viral supernatants were collected 48 h post-transfection. When mESCs or hESCs reached 60% confluence, lentivirus was added to the culture medium with 8 μg/mL polybrene. Forty-eight hours after infection, stably infected cells were selected using 2 μg/mL puromycin and/or 10 μg/mL blasticidin-S HCl.

### Immunoprecipitation-liquid chromatography-tandem mass spectrometry (IP-LC-MS/MS) analysis

The FLAG-tagged mouse *Tfcp2l1* gene was cloned into an empty vector to generate the EV-*Tfcp2l1* plasmid, which was then transfected into mouse embryonic stem cells (mESCs). Cells transfected with the empty vector (EV) served as the control group. IP-LC-MS/MS analysis was performed by oebiotech (Shanghai, China). Candidate interacting proteins were preliminarily screened using the threshold of a Sequest HT score > 5.

### Colony formation assay

Cells were seeded at a density of 2000 cells per well in 6-well plates and cultured for two weeks. Cell colonies were fixed with 4% paraformaldehyde and stained with 0.5% crystal violet (C8470, Solarbio) for 30 min. After three washes with PBS, images were captured, and colony numbers were quantified using ImageJ software.

### Alkaline phosphatase staining

Cells were fixed in 4% paraformaldehyde at room temperature for 2 min. After PBS washing, cells were incubated with alkaline phosphatase (AP) staining solution (C3206, Beyotime Biotechnology, China) for 30 min at room temperature in the dark. Following three PBS washes, cells were observed under a Leica DMI8 microscope.

### Cell proliferation assay

Cells were seeded into 96-well plates (Cat. 11510, Labselect, China) at a density of 2 × 10³ cells per well and incubated at 37 °C for 72 h. At each designated time point, the culture medium was discarded, and 100 μL of fresh culture medium was added to each well. Subsequently, 10 μL of CCK-8 solution (Cat. C0038; Beyotime, China) was added for further incubation for 1 h. The optical density (OD) value at 450 nm was measured using a microplate reader (SpectraMax M5; Molecular Devices, USA).

### Reverse-Transcription PCR (RT-PCR)

Total RNA was extracted from mESCs/hESCs with TRIzol reagent, followed by DNase I digestion to eliminate genomic DNA. Reverse transcription was carried out to generate cDNA. RT-PCR amplification was performed with gene-specific primers. PCR products were separated by agarose gel electrophoresis and visualized under ultraviolet light. Band intensities were quantified using ImageJ software. *Gapdh* was used as the loading control. Primer sequences are summarized in Table S3.

### Real-Time-quantitative PCR (RT-qPCR)

Total RNA was isolated using the total RNA extraction kit (Yeasen Biotechnology, China). Complementary DNA (cDNA) was synthesized using the cDNA Synthesis SuperMix for qPCR kit (Yeasen Biotechnology, China). RT-qPCR was performed with Universal SYBR qPCR Master Mix (11201ES03, Yeasen Biotechnology, China). Relative gene expression was calculated using the 2-DCq method and normalized to *β-actin*. Primer sequences are summarized in Table S4.

### Western blot

Total cellular protein was extracted by lysing cells in ice-cold RIPA lysis buffer (P0013B, Beyotime Biotechnology) supplemented with protease inhibitor cocktail. Chromatin-bound protein was extracted using the Nuclear and Cytoplasmic Extraction Kit (E101-01, Vazyme, China). Equal amounts of protein samples were separated by 6%, 10% or 15% SDS-PAGE and electro-transferred onto PVDF membranes. Membranes were blocked with 5% skim milk for 4 h and incubated with primary antibodies overnight at 4 °C. The primary antibodies used were as follows: TFCP2L1 (AF5726, R&D Systems, 1:1000), THRAP3 (85093-3-RR, Proteintech, 1:1000), BCLAF1 (HA722374, HUABIO, 1:1000), BCL2 (HR60235, ZENBIO, 1:1000), Caspase 3 (19677-1-AP, Proteintech, 1:1000), BAX (R22708, ZENBIO, 1:1000), γ-H2AX (ET1602-2, HUABIO, 1:1000), JMJD1C (A20153, ABclonal, 1:1000), MDC1 (HA724359, HUABIO, 1:1000), BRCA1 (A0212, ABclonal, 1:1000), RAP80 (EM1902-05, HUABIO, 1:1000), Pan Dimethyl-lysine (A5870, ABclonal, 1:1000), RNF8 (R381564, ZENBIO, 1:1000), ATM (A5908, ABclonal, 1:1000), p-ATM (S1981) (AP1030, ABclonal, 1:1000), ATR (A21253, ABclonal, 1:1000), p-ATR (S428) (AP1582, ABclonal, 1:1000), 53BP1 (A5757SP, ABclonal, 1:500), FLAG (F1804, Sigma, 1:1000), HA (51064-2-AP, Proteintech, 1:1000), OCT4 (11263-1-AP, Proteintech, China, 1:1000), KLF4 (R382455, ZENBIO, 1:1000), GATA4 (36966, Cell Signaling Technology, 1:1000), GATA6 (5851, Cell Signaling Technology, 1:1000), H3 (17168-1-AP, Proteintech, 1:1000), β-Actin (380624, ZENBIO, 1:1000), β-Tubulin (66219-1-Ig, ZENBIO, 1:1000). Membranes were incubated with secondary antibodies at room temperature for 2 h. Immunoreactive bands were visualized using a Tanon 5200multi gel imaging system (Shanghai, China).

### Immunofluorescence staining

Cells were washed three times with PBS and fixed with 4% paraformaldehyde for 30 min, followed by three additional PBS washes. Primary antibodies (1:500 dilution) were diluted in blocking buffer containing 5% BSA and 0.2% Triton X-100, and incubated with cells overnight at 4 °C. After three PBS washes, cells were incubated with species-specific fluorescent secondary antibodies (1:1000) and Hoechst 33342 (H3570, Invitrogen, 1:10000) diluted in blocking buffer for 1 h at 37 °C. After three PBS washes, images were acquired under a Leica DMI8 microscope. The primary antibodies used included TFCP2L1, THRAP3, SOX2 (66411-1-Ig, Proteintech, 1:500), 53BP1, RAD51 (A266856, ABclonal, 1:500), γ-H2AX, RNF8, MDC1, BRCA1 and RAP80.

### Embryoid-body formation assay

A total of 1 × 10⁷ mESCs or hESCs were cultured in suspension on ultra-low-attachment plates (3473, Corning) using standard serum-containing embryonic stem cell medium without LIF. Aggregates were cultured for up to 8 days. Samples were collected at designated time-points, lysed for RNA extraction and subjected to RT-qPCR analysis.

### Co-Immunoprecipitation (Co-IP) assay

Twenty microliters of Protein A+G Agarose (P2055, Beyotime Biotechnology) was incubated with 5 μL of TFCP2L1, THRAP3, FLAG, HA, JMJD1C, MDC1 or Pan DiMethyl-lysine antibody in 10% BSA blocking buffer overnight, and then washed three times with pre-cooled PBS. Cells were placed on ice, washed three times with pre-cooled PBS supplemented with protease inhibitors, and 600 μL IP lysis buffer was added. Cells were lysed with shaking at 4 °C for 15 min, scraped, and further incubated on a rotator at 4 °C for 15 min. Lysates were centrifuged at 12000 rpm for 15 min at 4 °C. A volume of 120 μL supernatant was reserved as the input sample. The remaining supernatant was incubated with pre-blocked 20 μL Protein A+G Agarose overnight at 4 °C. The resulting protein samples were analyzed by Western blot.

### ChIP-RT-qPCR

Chromatin immunoprecipitation (ChIP) was performed using the ChIP Assay Kit (P2078, Beyotime Biotechnology) according to the manufacturer’s instructions. Immunoprecipitation was performed using antibodies against TFCP2L1, THRAP3, RNA Pol II and RNA Pol II-S2P; IgG was used as the negative control. Enrichment levels were quantified by RT-qPCR. Primer sequences and binding-site information are provided in Table S5-S8.

### RIP-RT-qPCR

RNA immunoprecipitation (RIP) was carried out using the Pierce™ Magnetic RNA-Protein Pull-Down Kit (20164; Thermo Scientific) following the manufacturer’s protocol. Immunoprecipitation was performed with anti-TFCP2L1 and anti-THRAP3 antibodies; IgG served as the negative control. The abundance of co-precipitated RNA was measured by RT-qPCR. Primer sequences are listed in Supplementary Table S9.

### Luciferase assay

The promoter sequences of *Esrrb* and *Jmjd1c* (-2000 to +1) were inserted into the pGL6 plasmid to generate pGL6-*Esrrb* and pGL6-*Jmjd1c* reporter plasmids. Each reporter plasmid was co-transfected into embryonic stem cells together with Renilla luciferase control plasmid, *Tfcp2l1*-overexpression construct, and *Thrap3*-knockdown construct. Luciferase activity was measured 48 h post-transfection using the TransDetect® Double-Luciferase Reporter Assay Kit (FR201, Trans-Gen Biotech, China).

### mRNA-sequencing

Total mRNA was isolated from mESCs transfected with *Scramble* empty-vector, *Tfcp2l1* sh#1 and *Thrap3* sh#1 using TRIzol reagent (Invitrogen). Three biological replicates were prepared for each group. Mrna-sequencing was performed by oebiotech (Shanghai, China). Differentially expressed mRNAs were analyzed with the threshold of |log₂FC| > 1.0 and *p* < 0.05. The sequencing raw data have been deposited in the Gene Expression Omnibus (GEO) database.

rMATS (replicate multivariate analysis of transcript splicing) was utilized to analyze five categories of alternative splicing events: exon skipping (SE), alternative 5′ splice site (A5SS), alternative 3′ splice site (A3SS), mutually exclusive exons (MXE), and intron retention (RI). The difference in exon-inclusion level (delta “percent spliced in”, ΔPSI) between knockdown and control samples was used to quantify alterations in alternative-splicing events upon gene depletion. Alternative-splicing events with FDR < 0.05 and |ΔPSI| > 0.2 were regarded as statistically significant. The correlation of differential alternative-splicing events was calculated by Spearman correlation analysis. The Spearman’s correlation coefficient (rs) and corresponding *p*-values were computed using the “cor.test” function in the R statistical environment. Representative splicing events were visualized via rmats2sashimiplot (https://github.com/Xinglab/rmats2sashimiplot).

### Statistical Analysis

The number of biological replicates is indicated in each figure legend. All data are presented as mean ± SD. GraphPad Prism 8 software was used for data visualization. Unpaired two-tailed Student’s *t*-test was used for comparisons between two groups. One-way or two-way ANOVA followed by Sidak’s multiple comparisons test was applied for multiple-group comparisons. A *p*-value < 0.05 was considered statistically significant.

## Acknowledgements

This work was supported by the National Natural Science Foundation of China (32300606, 32270847).

## Conflict of Interest

The authors declare no competing interests.

## Author Contributions

Wei Lu: Investigation, Methodology. Xiaoxiao Wang: Validation, Investigation. Yuexuan Li: Validation, Formal analysis. Peng Chen and Yuqing Zhu, Formal analysis. Shou-Dong Ye: Supervision, Writing – Methodology, review & editing.

## Data Availability Statement

The data that support the findings of this study are available from the corresponding author upon reasonable request.

